# Structural characterization of LsrK to target quorum sensing and comparison between X-ray and homology model

**DOI:** 10.1101/2020.09.03.281394

**Authors:** Prasanthi Medarametla, Tuomo Laitinen, Antti Poso

## Abstract

Quorum sensing is being investigated as an alternative therapeutic strategy in antibacterial drug discovery to combat bacterial resistance. LsrK is an autoinducer-2 kinase, playing a key role in the phosphorylation of autoinducer-2 (AI-2) signalling molecules involved in quorum sensing. Inhibiting LsrK could result in reduced pathogenicity by interfering with the quorum sensing signalling. Previously, we have generated homology models to identify LsrK inhibitors using structure-based virtual screening and successfully found the first class of LsrK inhibitors. While conducting these studies, the crystal structure of LsrK was released providing us an opportunity to inspect the reliability and quality of our models. Structural analysis of crystal structure and homology models revealed the consistencies of constructed models with crystal structure in the structural fold and binding site. Further, binding characteristics and conformational changes are investigated using molecular dynamics. These simulations provided us insights into the protein function and flexibility that need to be considered during the structure-based drug design studies targeting LsrK.

## INTRODUCTION

Increased bacterial resistance has become a global health threat that urges the development of novel therapeutics. One of the main reasons for antibiotic resistance is the conventional mechanism of action of existing drugs i.e. targeting the protein synthesis, cell wall synthesis, and DNA replication^1,2^. Currently, novel strategies such as targeting virulence are prominent in antibacterial drug discovery^3,4,5^. Virulence strategies reduce the selection pressure on bacteria to develop resistance as these processes are not essential for bacterial growth^6,7,8^. The major focus of these strategies is on disrupting the host-pathogen interactions and reducing the pathogenicity by inhibiting the adhesion and toxin release, biofilm formation, and quorum sensing^9,10,11^. Quorum sensing (QS) is a process used by bacteria to communicate between the species and among the species. This communication controls the population-based behaviours and functions such as virulence factor secretion, biofilm formation, motility, bioluminescence, sporulation, and development of genetic competence^12,13^.

QS process is mediated by signalling molecules called autoinducers (AIs). These signalling molecules can be devided into three major groups: Acylated Homoserine Lactones (AHL), Autoinducer peptides (AIPs), and Autoinducer-2 (AI-2). AHLs are N-Acyl-L-homoserine lactones varying in their acyl chain length between 4 to 18 carbon atoms while AIPs are oligopeptides. Generally, AIPs are utilized by gram-positive bacteria whereas AHLs are used by gram-negative bacteria^14^. In contrast, AI-2 molecules are the universal signalling molecules that are used by both gram-positive and gram-negative bacteria. AI-2 produced in bacteria will be internalized from the extracellular environment into the cells by an ATP binding cassette (ABC) transporter system called the Lsr transporter. Further, AI-2 is phosphorylated by LsrK (encoded by the lsrK gene) inside the cell and undergoes further modifications by LsrF and LsrG. The isomerized Phospho-AI-2 is responsible for the lsr operon activation and inactivation of a repressor protein, LsrR^15^. Thus, hindering the phosphorylation of AI-2 can be a promising strategy in the design of antibacterial drugs. First reports by Zhu et al. provided details into the role of LsrK in AI-2 phosphorylation and its mechanism^16^. LsrK phosphorylates the AI-2 precursor, DPD (4,5-dihydroxy-2,3-Pentanedione) and thus regulates the AI-2 signalling and QS process. Implying the role of LsrK in QS signalling and virulence regulation, we explored LsrK to identify anti-virulence agents by employing the homology modelling and virtual screening approaches^17^. Recently, LsrK crystal structure (*E. coli*) was published by Ha JH et al. with a phosphocarrier protein, HPr. These studies revealed the role of LsrK kinase activity and how its activity is modulated by HPr protein^18^. It provides an opportunity to evaluate the quality of our homology models and gain insights into the details of protein flexibility and inhibitor design targeting the LsrK.

In this study, the main focus is to inspect the quality of homology models using the crystal structure and gain insights into protein flexibility. Molecular dynamics simulations were employed to understand the protein conformational changes of LsrK using crystal structures and constructed homology models. The structural details provided an understanding of the LsrK structure and the conformational changes occuring during the ATP and substrate binding. These details are helpful to guide the structure-based inhibitor design targeting the LsrK kinase to interfere with the quorum sensing process.

## RESULTS AND DISCUSSIONS

### Comparison of homology models and crystal structure

The release of *E. coli* LsrK structure (*ec*LsrK), crystallized with HPr protein, raised the interest to inspect the structural quality of our homology models (*S. typhimurium:st*LsrK). Both sequences were aligned to inspect the sequence differences using the ClustalW alignment server and depicted in Figure S1 using ENDScript 3.0. The sequence identities between *ec*LsrK and *st*LsrK were 82.64%. The major variation was found in Domain I of residues 76-85, and in Domain II of residues 419-424 and 496-503. For the structural comparison, homology models were aligned with the X-ray crystal structures using the protein structure alignment. The X-ray crystal structure is in the open state and thus the open state homology model was correlated with the X-ray crystal structure (CS) containing ATP, and a cryoprotectant (hexane-1,6-diol) (PDB ID: 5YA1). Alignment of the model and CS revealed that secondary structural elements (structure helices, strands and loops) are in good agreement with the X-ray structure except helix (α13) of residues 326-337 (Figure 1 and see Supporting Information for numbering Figure S2). This region was predicted as a loop in the homology model whereas in the CS it is a helix and located in the vicinity of the ATP binding site. However, none of the residues in this helix are interacting with the ATP. Further, RMSD was checked between the homology model and CS. The overall backbone RMSD was 2.89Å and the RMSD for binding site residues was 0.97Å.

**Figure 1.**
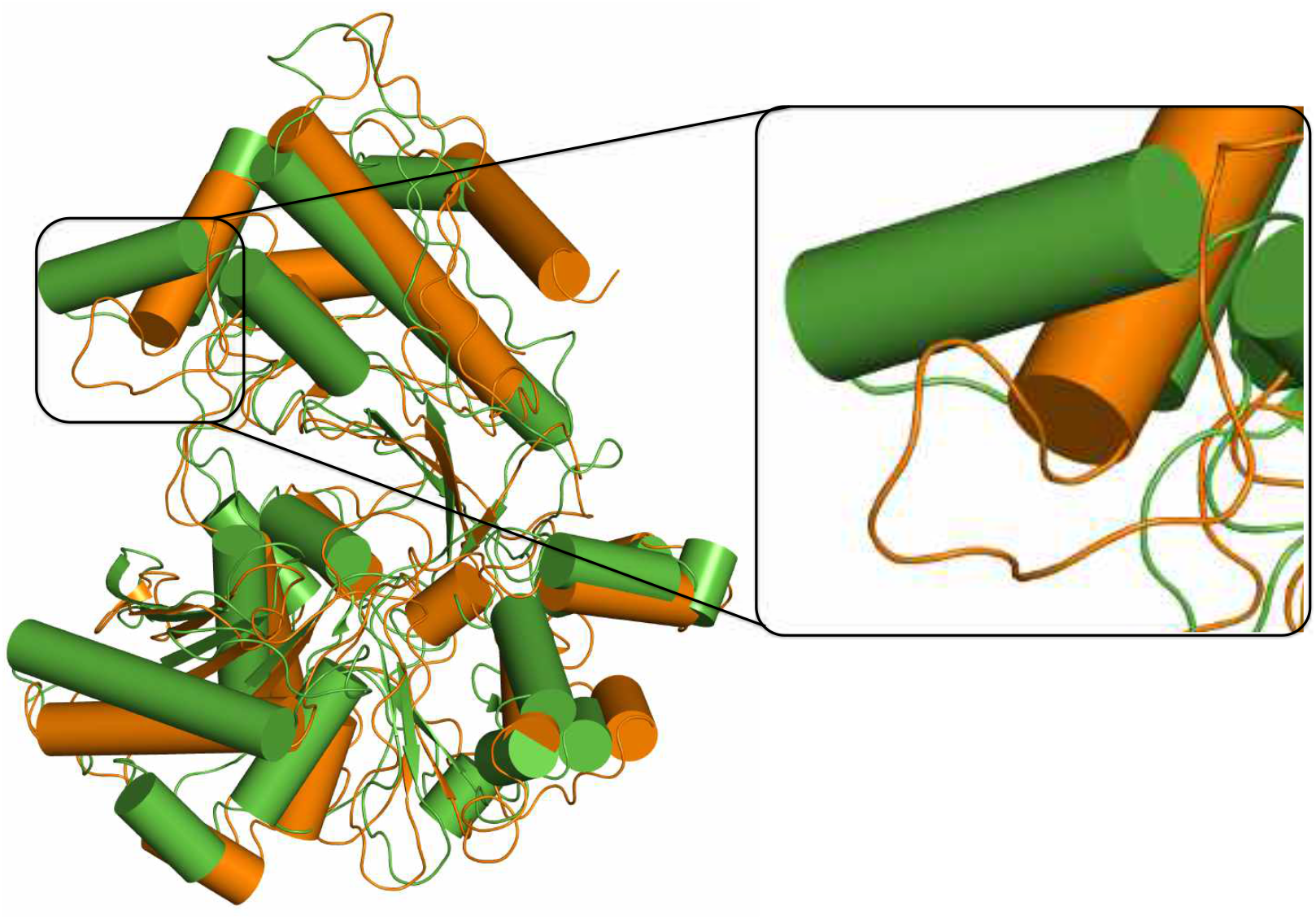
*Comparison of homology model (open state: orange colour) an1d x-ray crystal structure (5YA1: green colour). Secondary structural element* (α13) *predicted to be different in the homology model is highlighted using the box. Domain I and domain II are labelled*.

Further analysis revealed that CS and model are consistent in their location of the ATP binding site whereas they are different in the substrate location. The hexane-1,6-diol (cryoprotectant) present in the substrate binding site location is far from the modelled binding site (Figure SX). However, the binding site residues around the substrate binding site are similar in CS and homology model (Figure 2). To understand this in better detail, we inspected the X-ray structure and associated electron density maps. Density was not fully visible for the hexanediol (Figure SX). In addition, electron density was not identified for some of the regions (from 1-9, 46-54, 364-371 and 505-530) in the crystal structure^18^.

**Figure 2.**
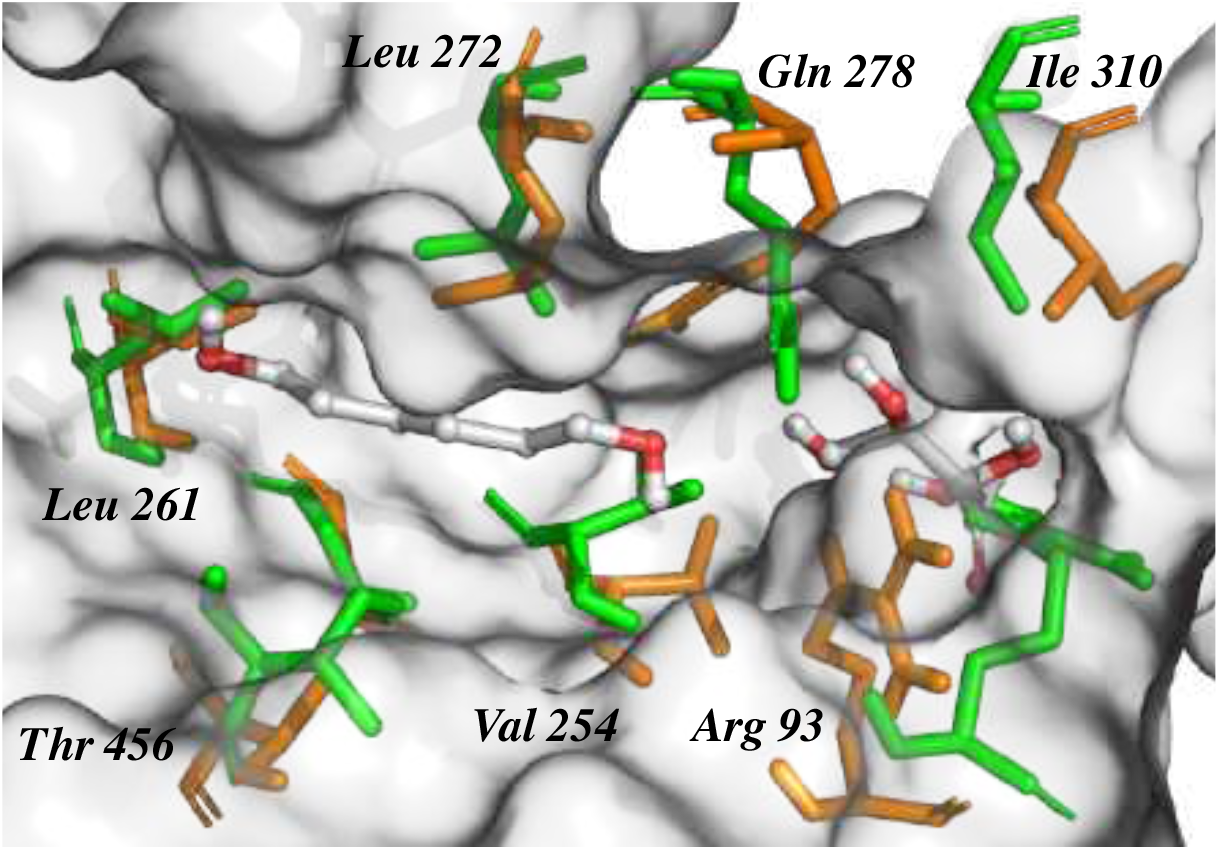
Binding site residues near the substrate binding site. Crystal structure is shown in green sticks and open state models in orange sticks.

### Protein flexibility and molecular dynamics

Crystal structure is available in apo form (CS-Apo), with ATP (CS-ATP), and with ADP (CS-ADP) in PDB IDs 5YA0, 5YA1 and 5YA2 respectively. There are no major conformational differences observed between the CS-Apo, CS-ATP, and CS-ADP. To understand how the protein flexibility and conformational changes occur during the substrate and ATP binding, molecular dynamic simulations were carried out on both homology models and all the crystal structures. Simulation trajectories were analysed for the Ca-atoms RMSD during the 500ns timescale (Figure 3).

**Figure 3.**
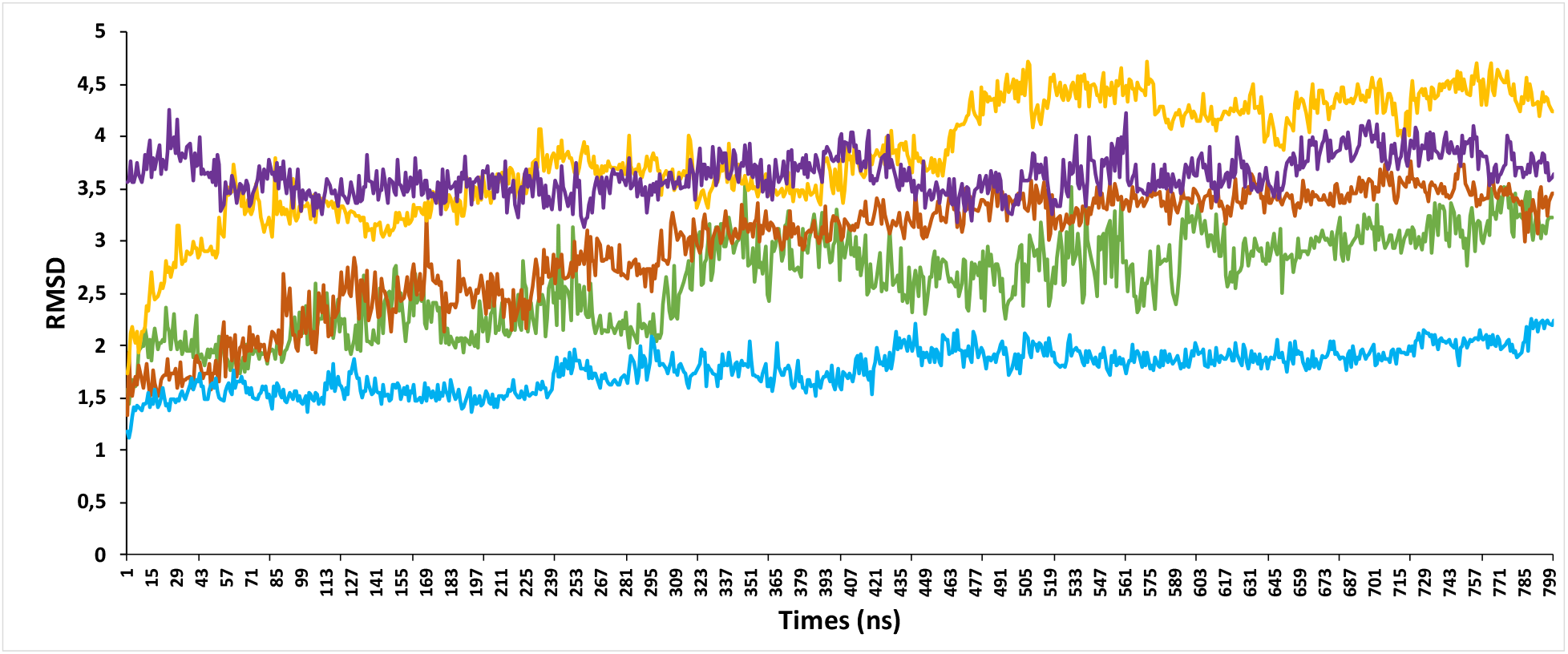
RMSD of Ca-atoms during the 500ns time scale of crystal structures: apo form (5YA0 or CS-Apo)-cyan lines, ATP bound form (5YA1 or CS-ATP)-green, ADP bound form (5YA2 or CS-ADP)-purple, open model-orange, and closed model-yellow.

RMSD of each trajectory is varying. The apo form (CS-Apo) is equilibrated at a less RMSD (1.5Å to 2Å) compared with other two forms and homology models. Open model (containing ATP and substrate) and CS-ATP were stabilized around 3Å. CS-ADP was stabilized at 3.5Å whereas closed model (including the substrate and ADP) was stabilized at the highest RMSD (around 4Å). Structural and mechanistic studies on FGGY carbohydrate kinase family members *E. coli* xylulose kinase (ecXK) and glycerol kinase (ecGK) revealed that phosphorylation occurs in the active state i.e. closed form. LsrK crystal structures (5YA0, 5YA1 and 5YA2) are reported to be in open conformation corresponding to the inactive state^18^. During the simulations, hexane-1,6-diol in the crystal structure (5YA1) and substrate in the open model (xylulose) were unstable in the binding site (supporting information Figures S3 and S4). This situation was contrasting in case of the closed model where the substrate was stable in the binding site during the 500ns timescale. This can be attributed to the interactions that occur between the ADP and substrate in the closed model during the simulation (Figure S4).

Further, simulation data was exploited to retrieve the information on protein flexibility information and protein structural movements in the protein. Root Mean Square Fluctuations (RMSF) values were used to identify flexible regions in the protein (Figure 4). Generally, N-terminal residues, C-terminal residues and loop regions show high fluctuations compared with other secondary structural elements such as helices and strands. During the 500ns simulation, the CS-ATP showed high fluctuations (RMSF > 5Å) at the loop regions of residues 35-47 (loop1) in domain I (located between α1 and β3) and 353-364 (loop2) in domain II (located between α14 and α15). The loops and major changes can be seen in the figure 4B. Minor fluctuations (RMSF <2Å) were noticed at residues 283-292 (loop3 between β12 and β13: the catalytic cleft). Loop1 and loop2 are in close proximity to the binding site region where the loop1 might be involved in the phosphorylation process (for protein numbering refer to Figure S2). Open model also showed high RMSF (> 5Å) in loop2 region and minor fluctuations (RMSF <3Å) in loop1 and loop 3. In addition to this, it is observed that the predicted loop (corresponding to helix α13 in CS) is also highly variable (RMSF > 5Å). To specifically investigate the highly dynamic regions and extreme movements in the protein structure principal component analysis was applied (Figure 5, Figure S5, and Figure S6).

**Figure 4.**
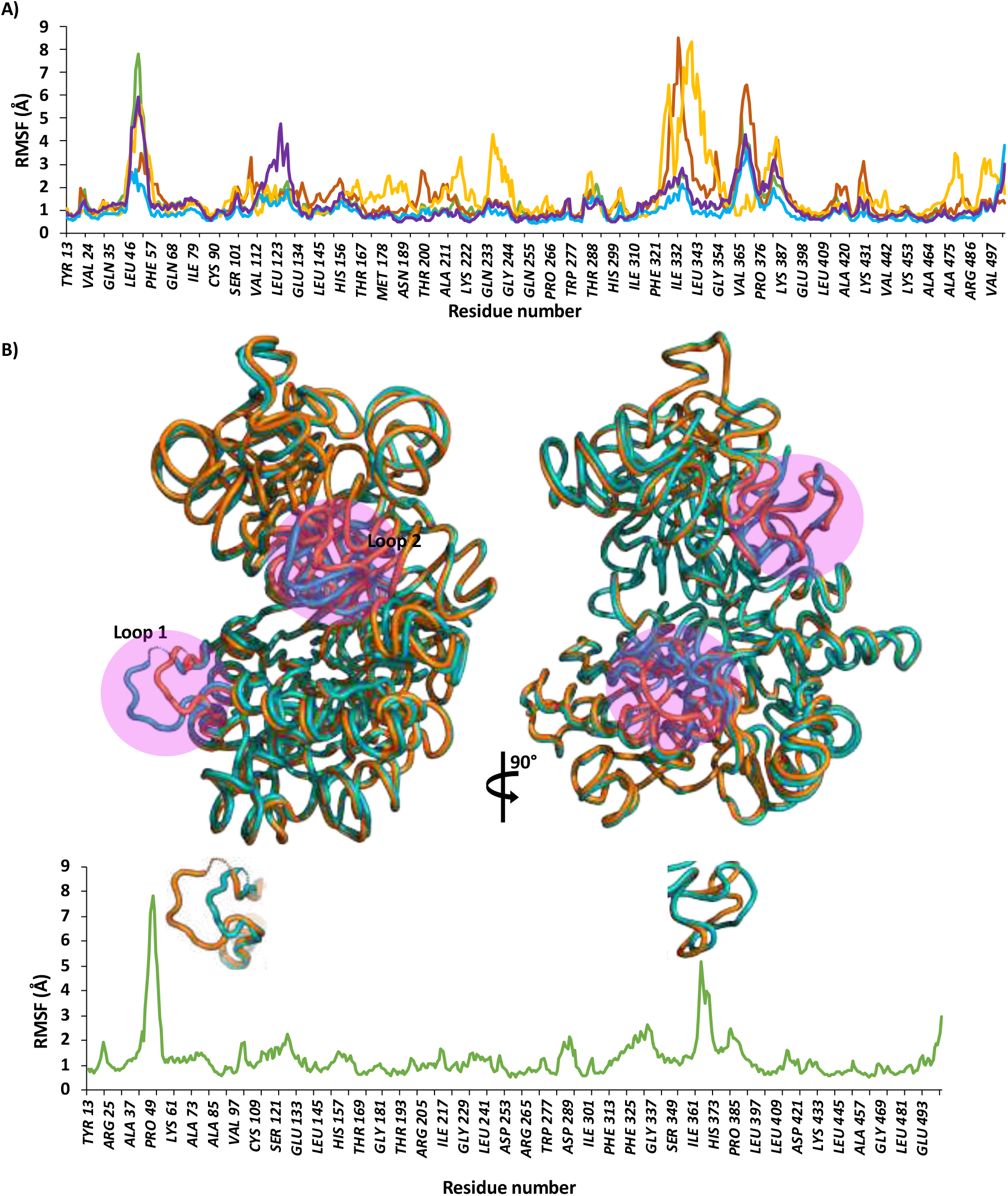
A) RMSF of Ca-atoms during the 500ns time scale. CS-Apo: cyan lines, CS-ATP: green, CS-ADP: purple, open model-orange, and closed model-yellow. The major fluctuating regions in all structures are residues 37-55, 320-330, and 360-380. B) To emphasize the regions in detail CS-ATP is shown. RMSF and related fluctuating regions are highlighted in the magenta circles of and showed them along the graph lines. Magnified version can be seen in Figure 5.

**Figure 5.**
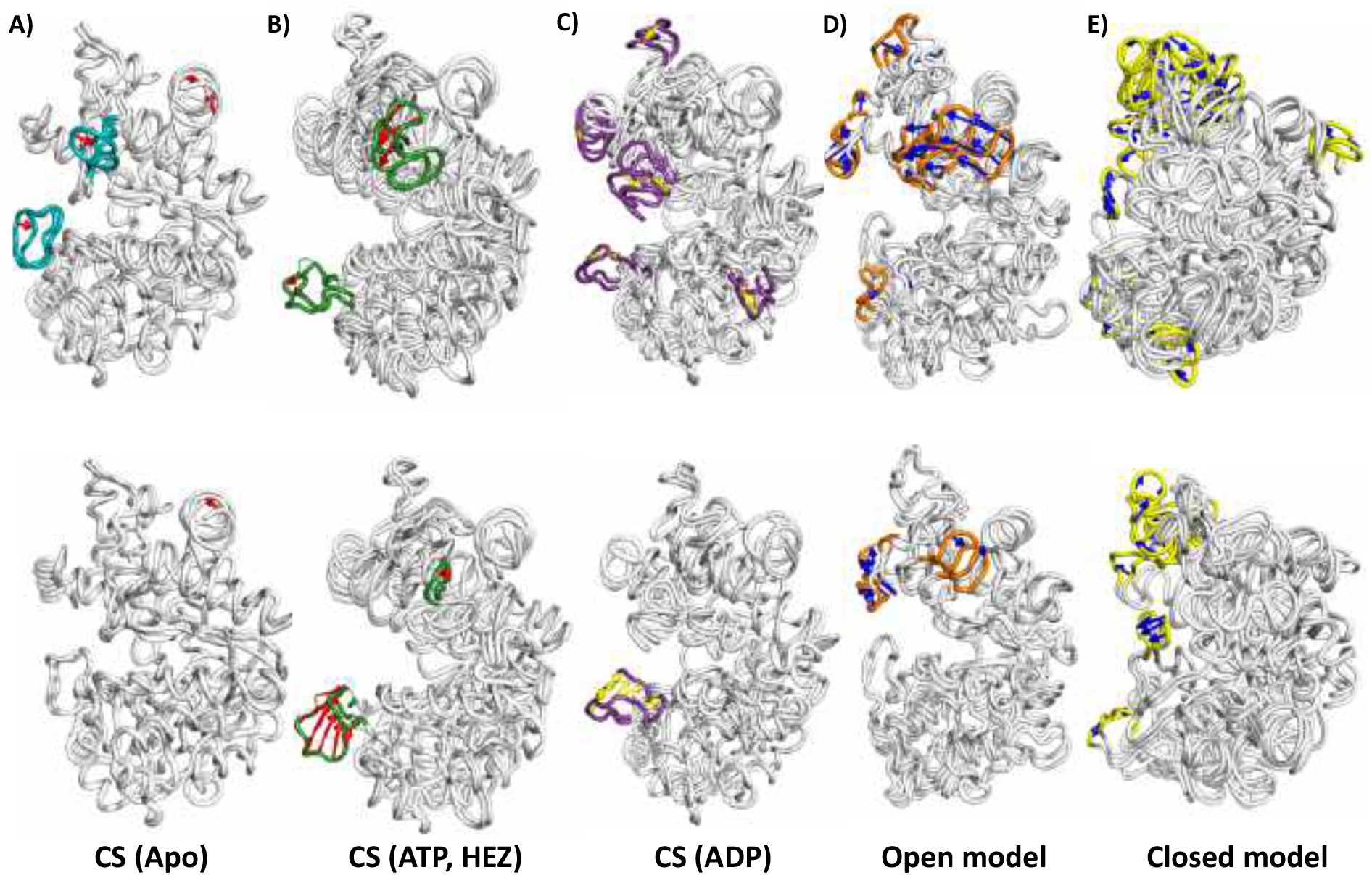
Highly variable regions in the crystal structure and homology models are presented using the PCA analysis: PC1 (top row) and PC2 (bottom row). Arrows (red and blue and yellow) indicate the direction of movement during the simulation. Extreme movements are highlighted using in coloured ribbons: teal (apo protein:5YA0), green (protein with ATP and hexane-1,6-diol:5YA1), purple (protein with ADP:5YA2), orange (open model with ATP and substrate), yellow (closed model).

As observed in the RMSF analysis, crystal structure (apo protein and ATP bound) showed large movements in loop regions i.e. loop1 of domain I and loop2 of domain II. These loop movements were also observed in the ADP bound structure (5YA2) and homology models. In addition to this, ADP bound LsrK showed large movements in the helix α14 (present near the ADP binding site) of domain II and α2 turn in domain I. Open model has also shown major movements near the helix α14 along with loop 1 and 2. During the simulation closed model showed extreme movements in domain II and small-scale movements in domain I (loop1 and loop proceeding α9). Overall, there are no structural movements observed near the substrate binding site in crystal structures and homology models. However, the loop1 (residues 35-45) stay in the proximity of catalytic cleft affecting the binding site size. This might be playing the role of gatekeeper during the substrate binding and the catalytic reaction in LsrK. Unfortunately, this is not established yet and this loop was not solved in the crystal structure (hence modelled using the Prime loop modelling for the simulations). Further, this analysis also revealed domain movements during the simulation and is also evident from the pocket shape and size analysis discussed in the next section.

### Understanding the domain movements using pocket shape and size analysis

The simulation trajectory clustering was performed based on the backbone RMSD using the Schrodinger trajectory clustering tool. To investigate the pocket size and volume of the substrate binding site, resulted ten clusters (centroids) were used. These clusters were subjected to pocket parameter prediction using Sitemap in Schrodinger KNIME workflows. Sitemap predicts possible druggable pockets and associated parameters such as size, volume, Sitescore, Dscore, hydrophobicity and hydrophilicity. Here, single binding site i.e. substrate binding site is analysed to predict the pocket parameters throughout the trajectory. Based on the predicted pocket volume, clusters were generated for all structures (CS-Apo, CS-ATP, CS-ADP, open model and closed model). All pocket parameters are tabulated in Supporting Information (Table T1-T5).

Analysis of these Sitemap pocket predictions revealed that volume of the pocket in each structure is varying throughout the simulation. The pocket shape analysis revealed that the change is due to the movement of protein domains (domain I and domain II). Figure 6 shows the ATP bound LsrK pocket changes in volume from 190.70Å to 294.63Å in panel A and panel B respectively. Pocket changes resulted majorly due to movements of loop 1 and domain II movement towards domain I. The domain movements and the pocket volume (lowest to highest) changes are shown in figure 7 for all structures. The structural changes in LsrK apo form, CS-ATP, and CS-ADP are evident from the pocket analysis. Crystallography predictions based on the superposition of xylulose kinase and glycerol kinase with LsrK revealed that ATP bound form (Figure 7B) is in open state. During the simulation, pocket volume among the three forms (apo, CS-ATP, and CS-ADP) changed from 112.16Å to 316.93Å. This demonstrates that LsrK undergoes conformational changes to accommodate ATP and substrate during the phosphorylation that incurs binding site volume changes presented here. Open model also showed considerable pocket volume changes from 245.24Å to 410.22Å. Closed model clusters sitemap predictions showed considerable domain I movement leading to the closure of binding site. Figure 7 shows the close packing of both domains in the closed model in comparison with the other structures.

**Figure 6.**
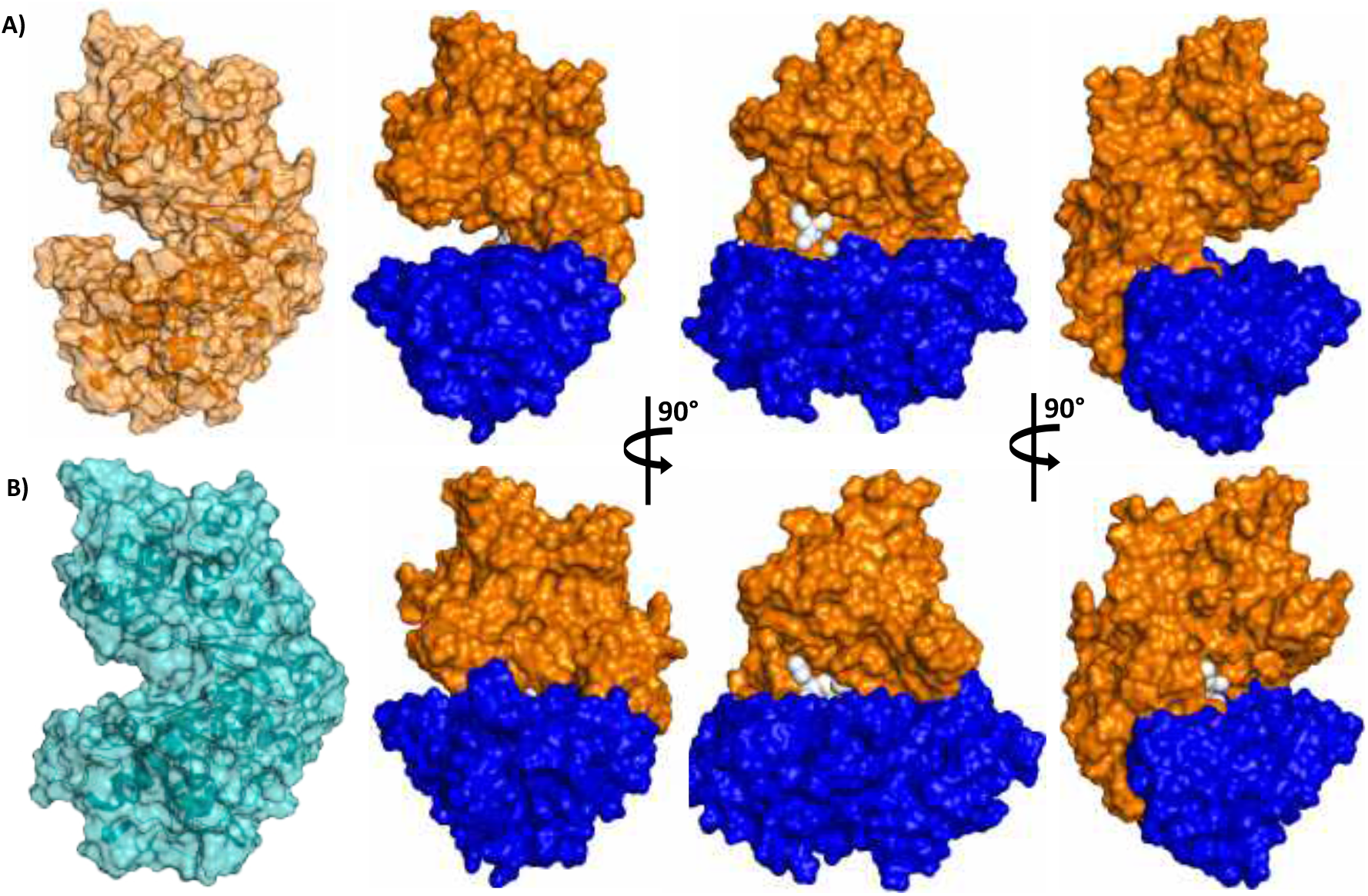
Pocket shape analysis of trajectory clusters of crystal structure (CS-ATP) showing the domain movements: domain I (blue) and domain II (orange). The snapshots of lowest pocket volume (A) and highest pocket volume (B) are visualized in orange and cyan colors. Other structures are shown in different angles to visualize the domain movements. Grey spheres indicate the site points.

**Figure 7.**
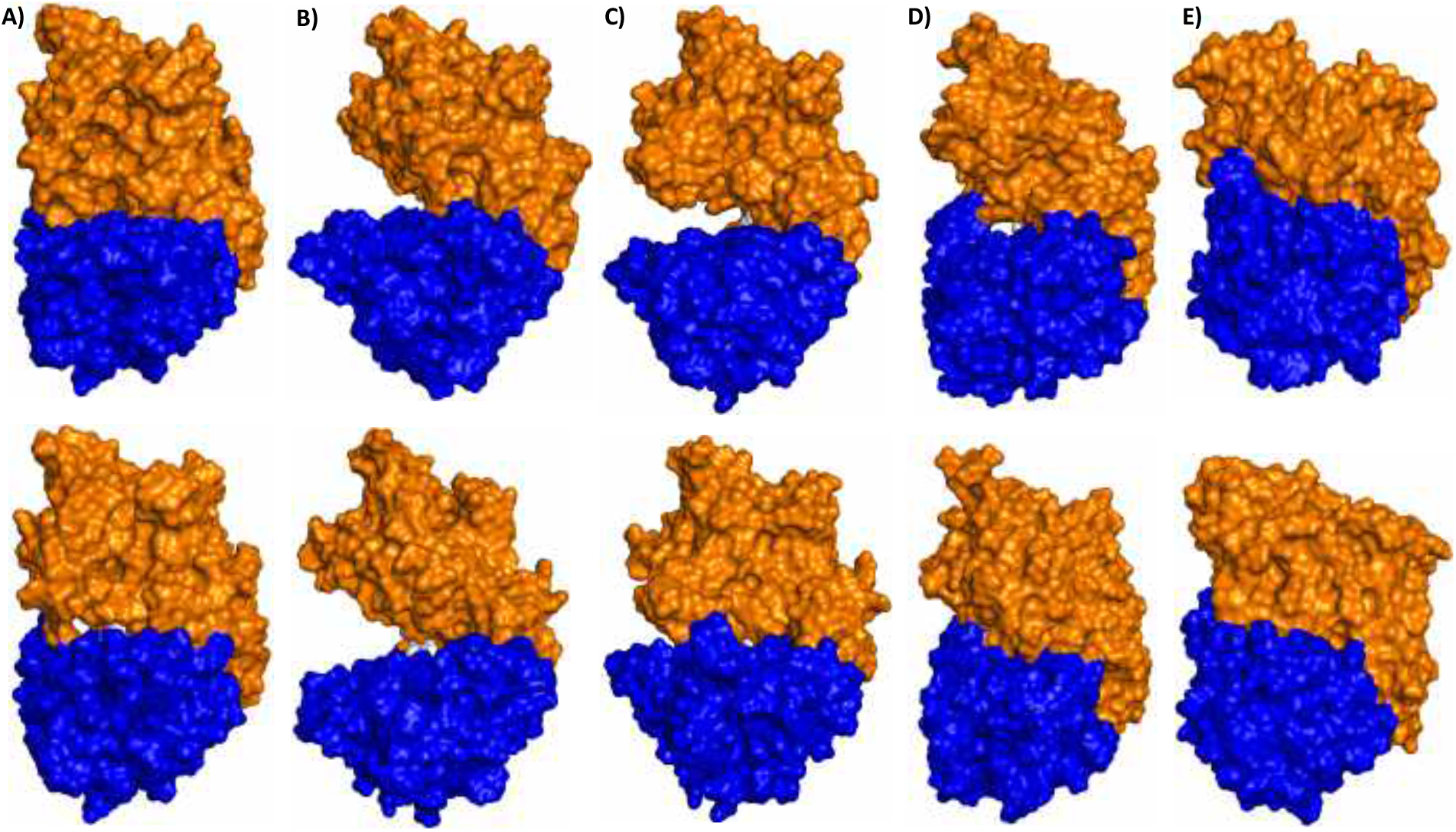
Trajectory clusters of crystal structure showing the domain movements: domain I (blue) and domain II (orange). Sitemap was utilized to predict the site volume. Top row represents the lowest site volume and bottom row with the highest site volume of apo structure (A), CS-ATP (B, CS-ADP (C), open model (D), and closed model (E). Grey spheres indicate the size of the pocket. Site volume and size are listed in supplementary information.

**Figure 8.**
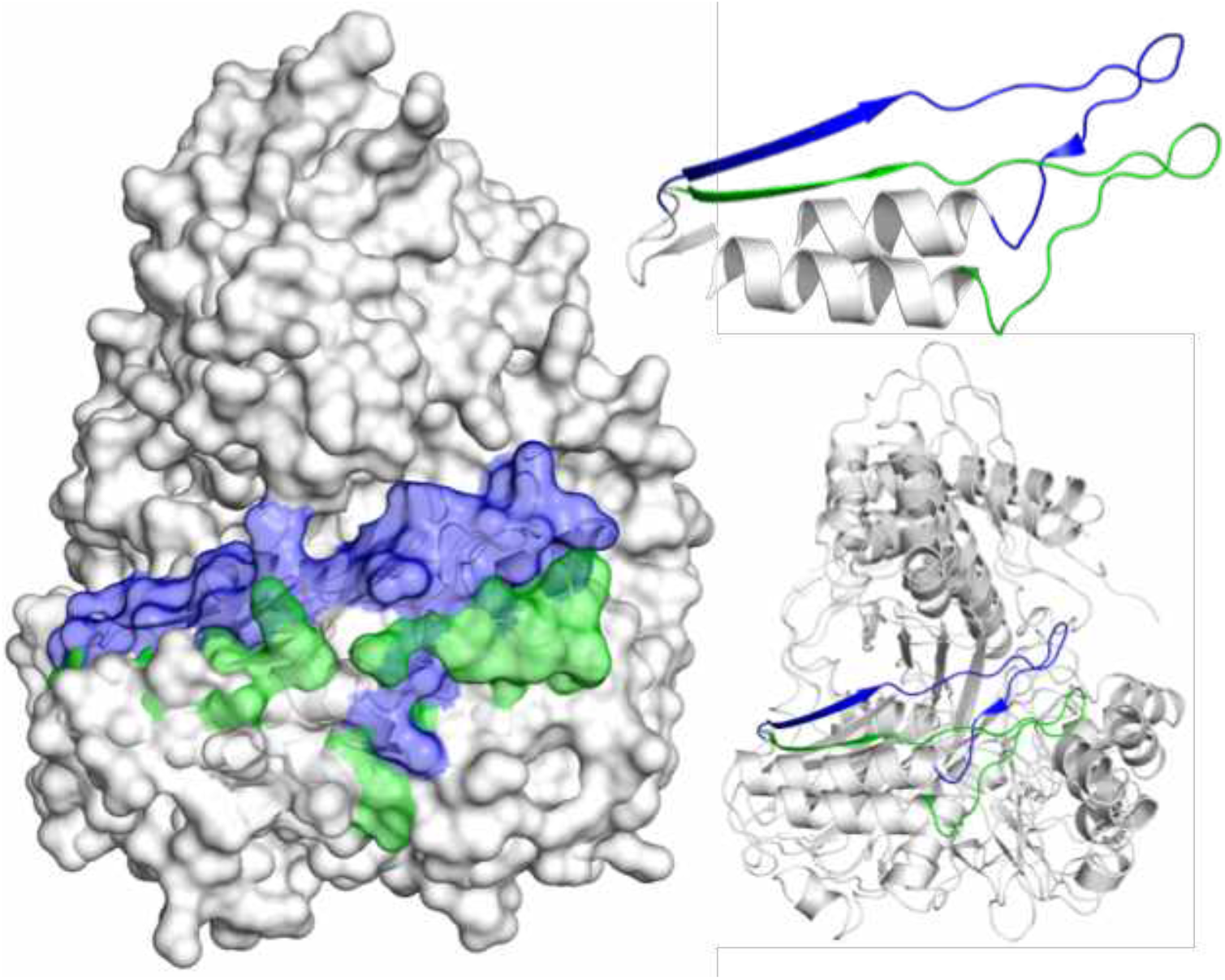
Superposition of xylulose kinase (PDBID:2ITM) and glycerol kinase (PDBID:1GLC). The conformational variations are highlighted in green color (xylulose kinase) and blue color (glycerol kinase).

To further confirm and investigate the conformational flexibility in related carbohydrate kinases, xylulose kinase and glycerol kinase structures were analysed. Large-scale differences were observed in domain I of crystal structures of xylulose kinase (PDB ID: 2ITM) and glycerol kinase (PDB ID: 1GLC). The domain I motion is in the vicinity of the beta sheet fold of substrate binding site (conserved hexokinase fold of the carbohydrate kinase family) and the loop 1. The major movements observed in LsrK during the dynamic simulations are coherent with the conformational flexibility observed in the xylulose kinase and glycerol kinase.

## CONCLUSIONS

The necessity to design new antibacterials has been raised to fight the emerging microbial resistance for the existing drugs and antibiotics. Alternative therapies such as targeting virulence to address the resistance is gaining interest in the research areas. One of such efforts is targeting the bacterial quorum sensing process that controls virulence and pathogenesis. LsrK is the key kinase involved in the quorum sensing process that regulates the *lsr* operon. Implying the role of LsrK in QS and virulence, computational methods are employed to understand the binding characteristics of LsrK and conformational changes associated with it. Our previous virtual screening driven by homology models led to the identification of first class of inhibitors for LsrK. The current study proves the quality of homology models and structural consistency with the crystal structure. Simulations are providing details about domain movements and structural flexibility that can help the structure-based drug design efforts to target LsrK binding site. Experimental studies are needed to depict the phosphorylation events occurring in the LsrK active site as this information would help LsrK targeted drug design further.

## METHODS

### Homology modelling

Structure of LsrK kinase of *Salmonella typhimurium* was modelled using FGGY carbohydrate kinase family proteins: xylulose kinase (23%) and glycerol kinase (26.5%) based on the sequence homology. Sequence alignment of LsrK and templates was carried out in prime and then manually edited with the help of multiple sequence alignment. Two models were built in Prime module of Schrödinger suite (Schrödinger release 2015-3: Prime, Schrödinger, LLC, New York, NY, 2015) using the templates in two different conformations. Template crystal structures *i.e*. xylulose kinase (3HZ6) and glycerol kinase (1GLC) were retrieved form PDB. Prime refinement protocol was used to predict the side chains and then minimized using OPLS_2005 force field^19^. Models were validated for their stereochemical quality and other parameters using tools such as Ramachandran Plot, ModVal and ERRAT factor. In Ramachandran plot, 98% and 94.8% residues are in allowed regions of open model and closed model. The ERRAT quality factor of the open model is 82.7 and closed model is 80.1. ModVal Predicted GA-341 is > 0.7 which indicates that models are reliable with ≥ 95% probability of the correct fold. Based on these statistical parameters, homology models were found to be of optimum quality that can be used for virtual screening purposes. Thus, homology models were further used to identify LsrK inhibitors using structure-based virtual screening. The detailed procedures of the homology modelling, virtual screening and experimental bioassays can be found in our previous paper^17^.

### Comparison of X-ray structure and Models

#### Sequence and Structure Analysis

LsrK sequences (*E. coli* and *S. typhimurium*) were retrieved from UniprotKB and analysed for the identity and similarity using ClustalW alignment server. Further, to inspect the structural differences crystal structures (5YA0, 5YA1, 5YA2) were downloaded from the RCSB PDB. Crystal structures often have problems such as lack of hydrogens, missing residues, incorrect protonation states etc. Hence, proteins are prepared using the protein preparation wizard (PPW) module in Schrödinger (Schrödinger Release 2019-3: Protein Preparation Wizard; Epik, Schrödinger, LLC, New York, NY, 2019) to assign bon orders, add hydrogens, fill missing side chains and loops, and generate ionization states to hereto atoms according to pH. Here, missing residues (45-55) were added based on the sequence using serial loop sampling method in Prime module (Schrödinger Release 2019-3: Prime, Schrödinger, LLC, New York, NY, 2019). Further, h-bond assignment was optimized and then structures were subjected to restrained minimization using OPLS3e force field^20^. To perform the structure analysis, prepared crystal structures and homology models were aligned using the Schrödinger superpose tool based on the backbone C-alpha atoms and calculated the RMSD. Further, structures were visually inspected near the ATP and substrate binding site to find the differences.

#### Molecular Dynamics

Crystal structures (PDB ID: 5YA0, 5YA1, 5YA2) and homology models (open and closed forms) were subjected 500 ns simulations using GPU-accelerated Desmond program using OPLS3e force field (Schrödinger Release 2019-3: Desmond, New York, NY, 2019). To carry out simulations, systems were built using system builder panel of Desmond. Systems were solvated with the T1P3P solvent model and orthorhombic periodic boundary conditions (of box size 10Å) and neutralized by placing ions of sodium (Na+). The solvated systems were equilibrated for 160 ps prior to the production runs using default NPT ensemble relaxation protocol in Desmond. This includes several stages: i) 100 ps of Brownian dynamics with NVT ensemble at 10 K temperature by posing restraints on solute heavy atoms ii)12 ps simulations with NVT ensemble using a Langevin thermostat (of 10 K) and restraints on solute heavy atoms iii)12 ps simulations of NPT ensemble using a Langevin thermostat (of 10 K), Langevin barostat (of 1 atm), and restraints on solute heavy atoms iv) solvating the pocket v)12 ps simulations with NPT ensemble using Langevin thermostat (of 300 K), Langevin barostat (of 1 atm), and restraints on solute heavy atoms vi) 24 ps simulations with NPT ensemble using Langevin thermostat (of 300 K), Langevin barostat (of 1 atm), and no restraints. Production runs were carried out for 500 ns using an NPT ensemble at 310 K with Nose-Hoover chain Langevin thermostat method and a pressure of 1.01 bar using the Martyna-Tobias-Klein barostat method. Initial atom velocities were assigned by randomization. The Coulombic interactions were truncated by using a cutoff value of 9 Å. RESPA based integration method was used with a 2.0 fs timestep and structures (around 1000 frames) were saved for every 100 ps for further analyses. Primary trajectory analyses were carried out using the simulation interaction diagram (SID) tool in Desmond. This tool calculates the RMSD and RMSF of the protein and ligand based on the selected atoms of the reference structure (i.e. RMSD and RMSF of each frame in the trajectory to the reference frame). It also provides information of protein-ligand interactions during the simulation and the ligand torsions. Here, RMSD and RMSF for all structures (5YA0, 5YA1, 5YA2, open model, and closed model) were calculated based on their Ca-atoms to the initial protein structure coordinates (X-ray structure and model). The RMSD and RMSF values were plotted using MS excel. Further, essential dynamics analysis was done using principal component analysis (PCA) using Bio3D^21^ in R console. PCA analysis was conducted with reference to the backbone Ca atoms of protein structures. Bio3D is used for the PCA analysis and results are visualized using Modevectors script^22^ in PyMOL 2.4.0 (The PyMOL Molecular Graphics System, Version 2.0 Schrödinger, LLC). Further, trajectory cluster analysis was performed using Desmond trajectory clustering tool to investigate the binding site volume changes. Each trajectory was subjected to this clustering tool to generate ten clusters representing the full trajectory. These ten clusters were further utilized to evaluate the substrate binding site using Sitemap module in Knime workflows^23^. The resulted structures were clustered based on binding site volume and visualized in PyMOL 2.4.0 (The PyMOL Molecular Graphics System, Version 2.0 Schrödinger, LLC.

## Supporting information

The additional files are available as pdf.

## ASSOCIATED CONTENT

### Supporting Information

The additional files are available as pdf.

## AUTHOR INFORMATION

### Author Contributions

The original draft was written by P.M. Manuscript was edited and reviewed by T.L and A.P. All authors have given approval to the final version of the manuscript.

## ACKNOWLEDGMENT

We also thank CSC - IT Center for Science Ltd. Finland for the use of their facilities, software licenses, computational resources and the Biocenter Finland/DDCB for financial support.

