## Supplementary material for "Structural characterization of LsrK to target quorum sensing and comparison between X-ray and homology model": The additional files are available as pdf.

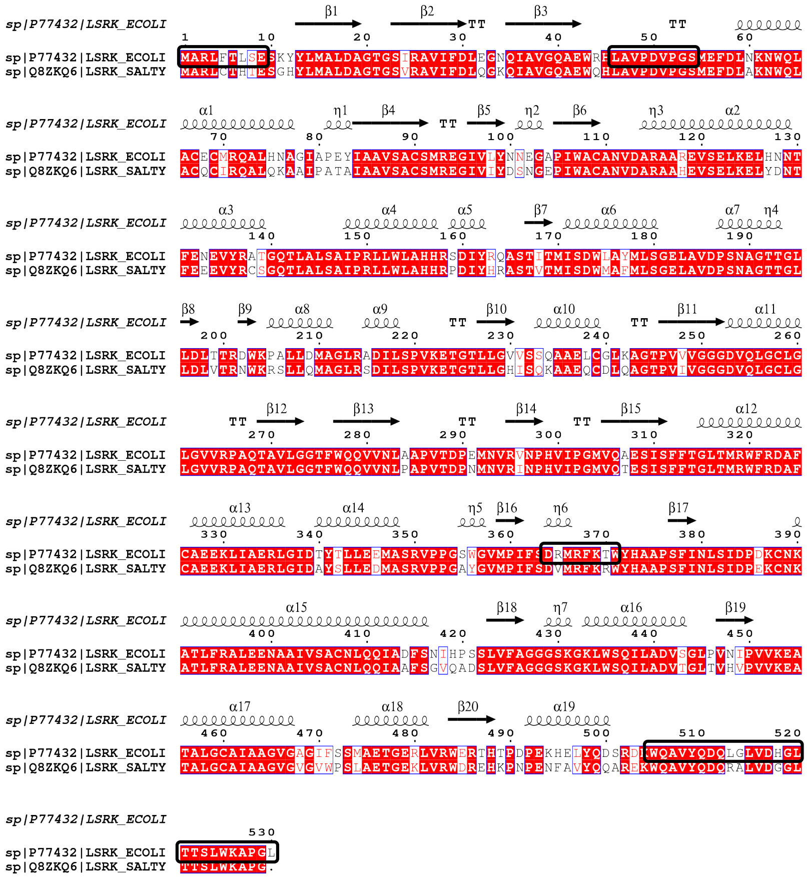

*Figure S1 Sequence alignment of E. coli LsrK kinase and S. typhimurium LsrK kinase. Red shading shows the identical residues. Secondary structural elements of the crystal structure were shown above the alignment. α-Helices were shown as squiggles and β-strands as arrows, strict β-turns as TT and strict α-turns as TTT. Regions that were not identified in the X-ray structure are highlighted with black boxes.*

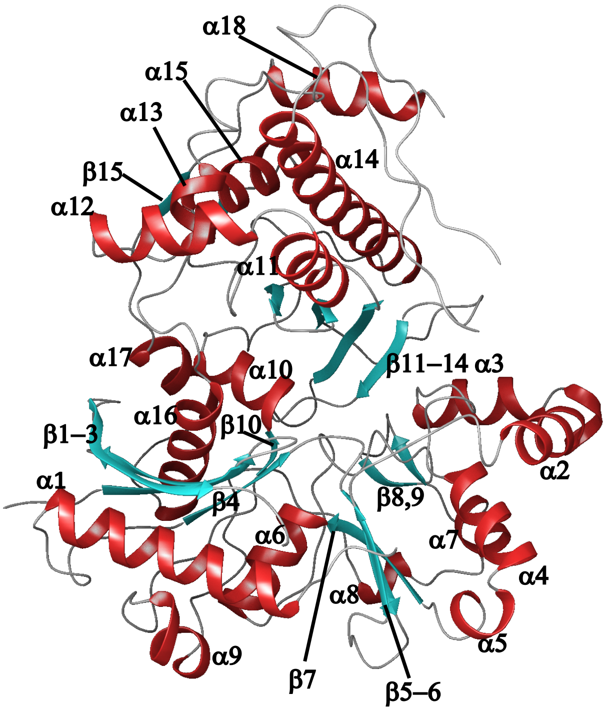


*Figure S2. Numbering for the reference of structure of LsrK (PDBID: 5YA1)*

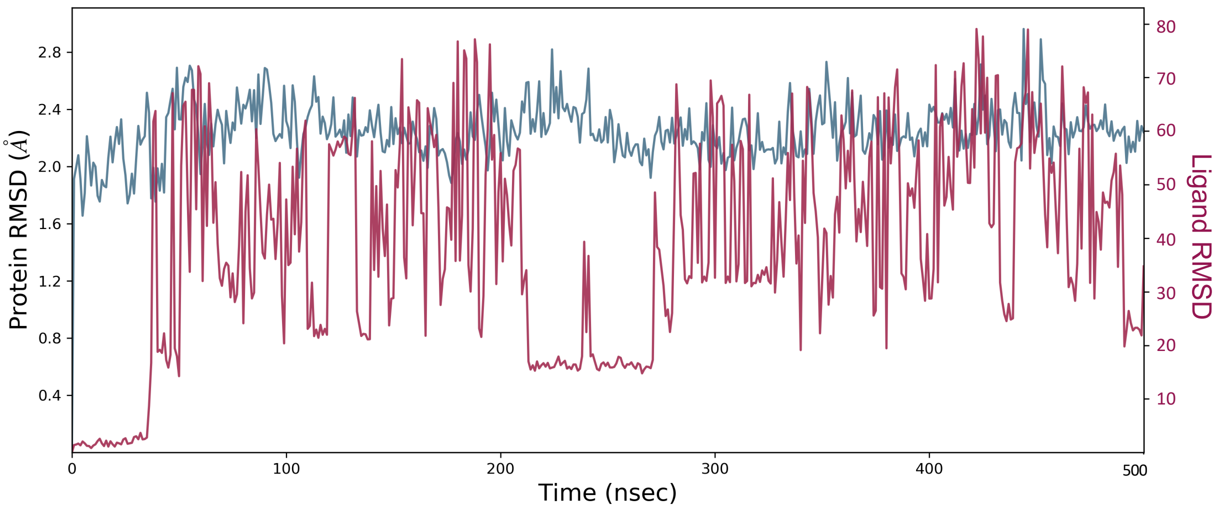


*Figure S3. Protein (Cα-atoms) RMSD and ligand RMSD of crystal structure (5YA1) during the dynamic simulations. RMSD is shown during the simulation timescale. Blue lines in the graph indicate protein RMSD and brown lines indicate ligand RMSD with respect to the protein.*

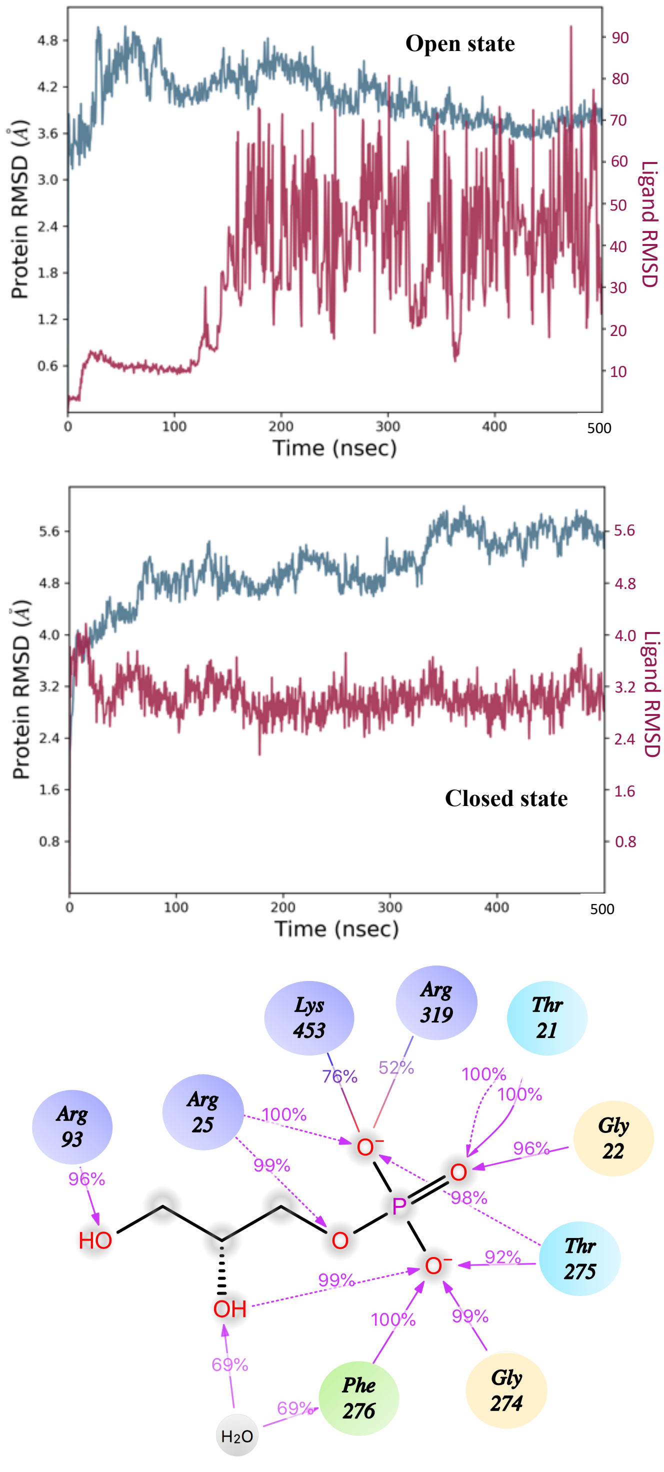


*Figure S4. RMSD of protein and ligand during the 500ns simulations of homology models. Protein-ligand interaction diagram of the closed model during the 500ns simulation time scale. Interactions and time of residence (%) are depicted.*

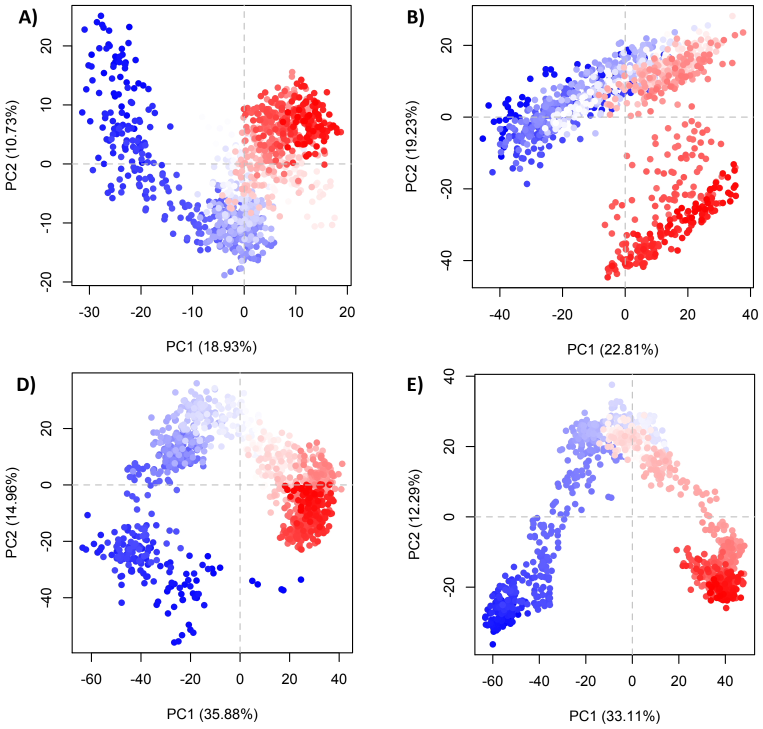


*Figure S5. Principal Component Analysis (PCA): two components PC1 (X-axis) and PC2 (Y-axis) are shown in the graph. A) Apo form (5YA0) B) CS-ATP bound form C) CS-ADP bound form D) Open model, and E) Closed model.*

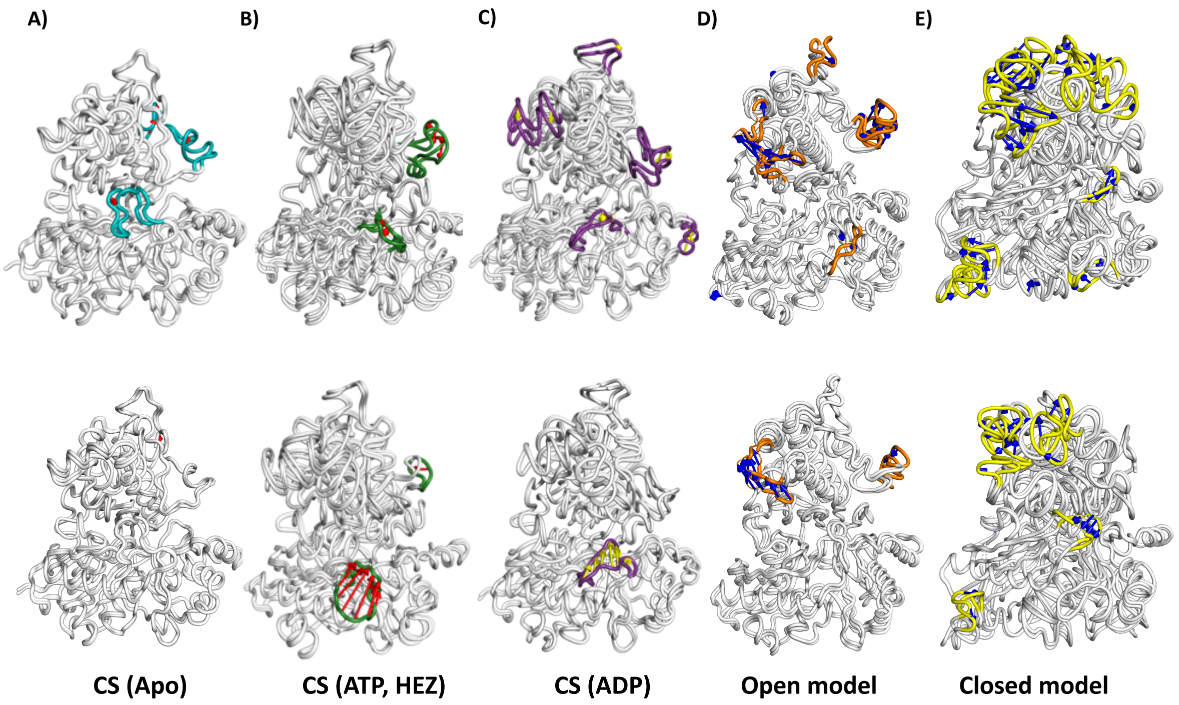


*Figure S6. Highly variable regions in the crystal structure and homology models are presented using the PCA analysis: PC1 (top row) and PC2 (bottom row). Arrows (red and blue and yellow) indicate the direction of movement during the simulation. Extreme movements are*

*highlighted using in coloured ribbons: teal (apo protein:5YA0), green (protein with ATP and substrate:5YA1), purple (protein with ADP:5YA2), orange (open model with ATP and substrate), yellow (closed model).*

*Table T1. Sitemap predicted pocket parameters for the trajectory cluster centroids of ATP and substrate bound crystal structure (5YA0).*

| Cluster Pocket Number | Size | Volume |
|---|---|---|
| Pocket 1 | 60 | 300.12 |
| Pocket 2 | 67 | 187.96 |
| Pocket 3 | 88 | 252.44 |
| Pocket 4 | 69 | 294.98 |
| Pocket 5 | 89 | 316.93 |

*Table T2. Sitemap predicted pocket parameters for the trajectory cluster centroids of ATP bound crystal structure (5YA1).*

| Cluster Pocket Number | Size | Volume |
|---|---|---|
| Pocket 1 | 40 | 190.70 |
| Pocket 2 | 35 | 146.80 |
| Pocket 3 | 51 | 243.53 |
| Pocket 4 | 55 | 294.637 |

*Table T3. Sitemap predicted pocket parameters for the trajectory cluster centroids of ADP bound crystal structure (5YA2).*

| Cluster Pocket Number | Size | Volume |
|---|---|---|
| Pocket 1 | 19 | 112.16 |
| Pocket 2 | 59 | 284.69 |
| Pocket 3 | 44 | 177.67 |
| Pocket 4 | 31 | 133.08 |
| Pocket 5 | 49 | 227.40 |

*Table T4. Sitemap predicted pocket parameters for the trajectory cluster centroids of Open model.*

| Cluster Pocket Number | Size | Volume |
|---|---|---|
| Pocket 1 | 31 | 245.24 |
| Pocket 2 | 66 | 297.72 |
| Pocket 3 | 87 | 332.71 |
| Pocket 4 | 109 | 380.73 |
| Pocket 5 | 98 | 410.22 |

*Table T5. Sitemap predicted pocket parameters for the trajectory cluster centroids of Closed model.*

| Cluster Pocket Number | Size | Volume |
|---|---|---|
| Pocket 1 | 84 | 132.05 |
| Pocket 2 | 158 | 227.75 |
| Pocket 3 | 145 | 109.57 |
| Pocket 4 | 156 | 261.70 |
